# A pipeline to detect the relationship between transposable elements and adjacent genes in host genomes

**DOI:** 10.1101/2021.02.25.432867

**Authors:** Meguerditchian Caroline, Ergun Ayse, Decroocq Veronique, Lefebvre Marie, Bui Quynh-Trang

**Author notes:** **Cite as**: Meguerditchian C, Ergun A, Decroocq V, Lefebvre M, Bui Q-T (2021) A pipeline to detect the, relationship between transposable, elements and adjacent genes in host, genomes. bioRxiv, 2021.02.25.432867, ver. 4 peer-reviewed and recommended by Peer community, in Genomics., https://doi.org/10.1101/2021.02.25.432867. This article has been peer-reviewed and recommended by *Peer Community in Genomics* https://doi.org/10.24072/pci.genomics.100010.

## Abstract

Understanding the relationship between transposable elements (TEs) and their closest positional genes in the host genome is a key point to explore their potential role in genome evolution. Transposable elements can regulate and affect gene expression not only because of their mobility within the genome but also because of their transcriptional activity. A comprehensive knowledge of structural organization between transposable elements and neighboring genes is important to study TE functional role in gene regulation. We implemented a pipeline to display the relationship between the distribution of TEs and adjacent genes in host genomes. Our tool is freely available here: https://github.com/marieBvr/TEs_genes_relationship_pipeline.

## Introduction

Transposable elements (TEs) were first discovered in maize by Barbara McClintock in 1948 [1]. They are mobile repetitive DNA sequences, and correspond to a significant part of eukaryotic genomes. These elements make up nearly half of the human genome [2], and about 85% of the maize genome [3]. There are many different types and structures of TEs [4], but they can be divided into two major classes: Class I contains elements called retrotransposons and Class II groups DNA transposon elements. These two classes are distinguished by their transposition mechanisms. The retrotransposons are firstly transcribed into RNA then retrotranscribed into double-stranded sequences to integrate into the genome, whereas the DNA transposons achieve their mobility based on an enzyme called transposase, which helps to catalyze DNA elements from its original location to insert into a new site within the genome. Due to their transposition and the act of transcription itself, the TEs can regulate gene expression by modifying genes into pseudogenes, by triggering alternative splicing or by modifying epigenetic marks [5], [6], [7], [8]. Experimental findings hint that TE insertions are targeted beyond the apparent mechanistic necessity of open chromatin [9]. Other examples have argued the possibility that more complex molecular mechanisms have evolved and made evident that TEs are an integral part of the regulatory toolkit of the genome [10], [11]. Studying the relationship between TEs and the surrounding genes will help determine the role of TEs in the many layers of gene expression regulation as well as their contribution to the plasticity of eukaryotic genomes and the evolution and adaptation of species. Many existing tools allow the users to predict TE locations in the genome, such as LTRpred [12], PiRATE [13] or REPET [14], [15]. Others tools can reveal the relationship between TEs and host sequences at transcriptome level. LIONS [16] is one of them that detects and quantifies transposable element initiated transcription from RNA-seq. Another software that can provide the screening and selection of potentially important genomic repeats is GREAM (Genomic Repeat Element Analyzer for Mammals) [17]. However, this web-server can offer the TE-gene neighborhood only within mammalian species. There are no tools for revealing positional relationship between TEs and host coding sequences that can be applied for any species. In order to help biologists in studying TE impacts upon gene expression, we developed a pipeline allowing users to report the association between TEs and their closest genes, such as its location and distance to adjacent genes within the genome. Our dedicated command line tool can be used with TE annotation from any prediction tool as well as any genome assembly and annotation. This pipeline also provides a graph visualization in R.

## Materials and methods

### 1. General workflow

Two input files are needed, a GFF file annotation of the host genome and a TSV file containing information about TE annotation. The Apricot dataset (*Prunus mandshurica*) and two different TE annotations were used to implement our pipeline: TE annotation from the REPET package and LTR retrotransposon annotation from LTRpred software. The gene and TE annotation files are firstly optimized for data analysis, by removing duplicate lines and sorting out by chromosome and by start position in ascending order. Then, the files are processed with Python in order to analyze the relationship between nearest genes. Our pipeline could detect and report all scenarios that could occur between TEs and the closest genes in the host genome as shown in the Figure 1. The Python output result is then used in R to create tables and graphs to visualize TE-gene relationships. The whole workflow is descripted in Figure 2.

**Figure 1.**
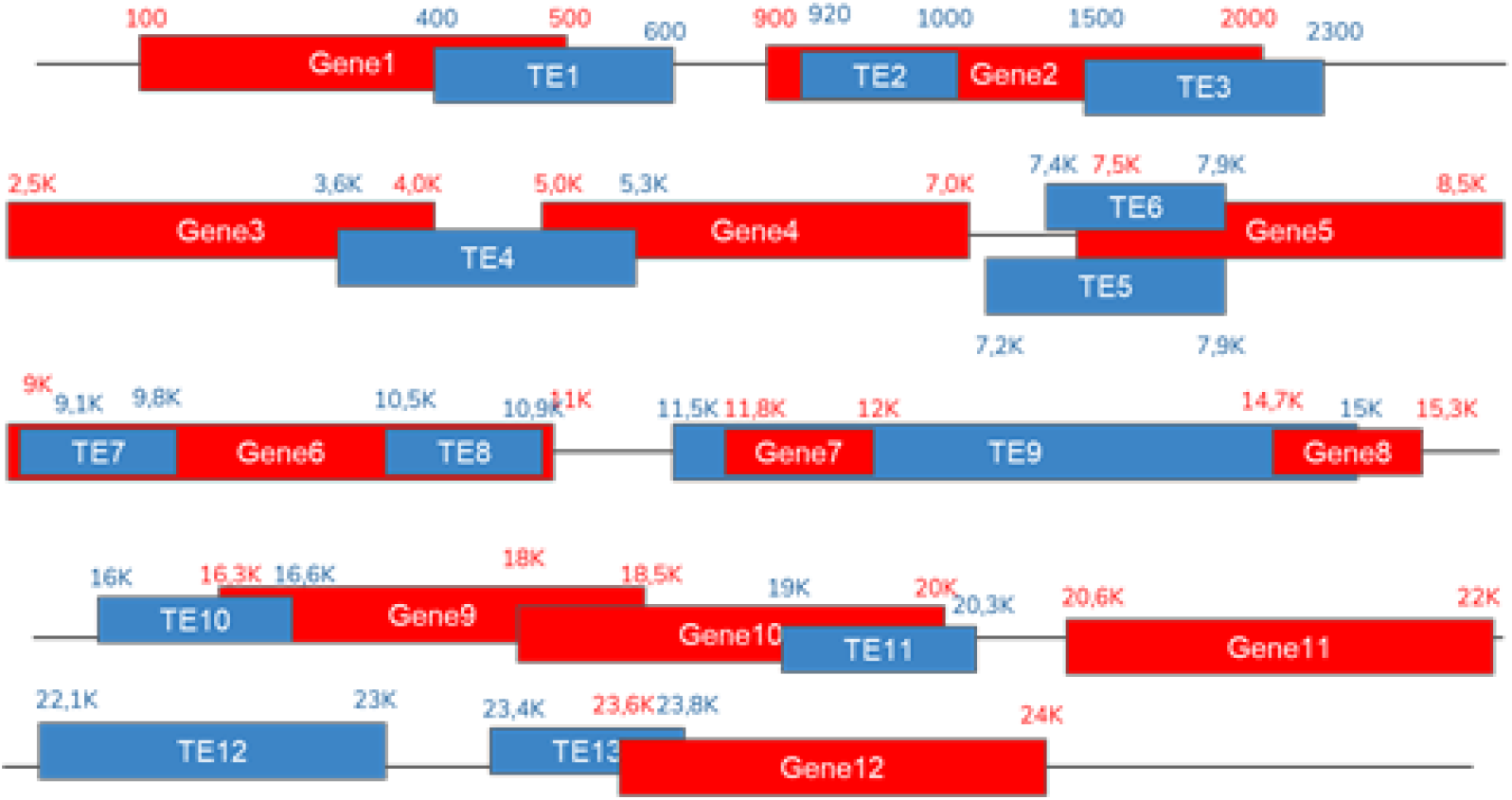
Scenarios of relationship between TEs and the closest genes in the host genome.

**Figure 2.**
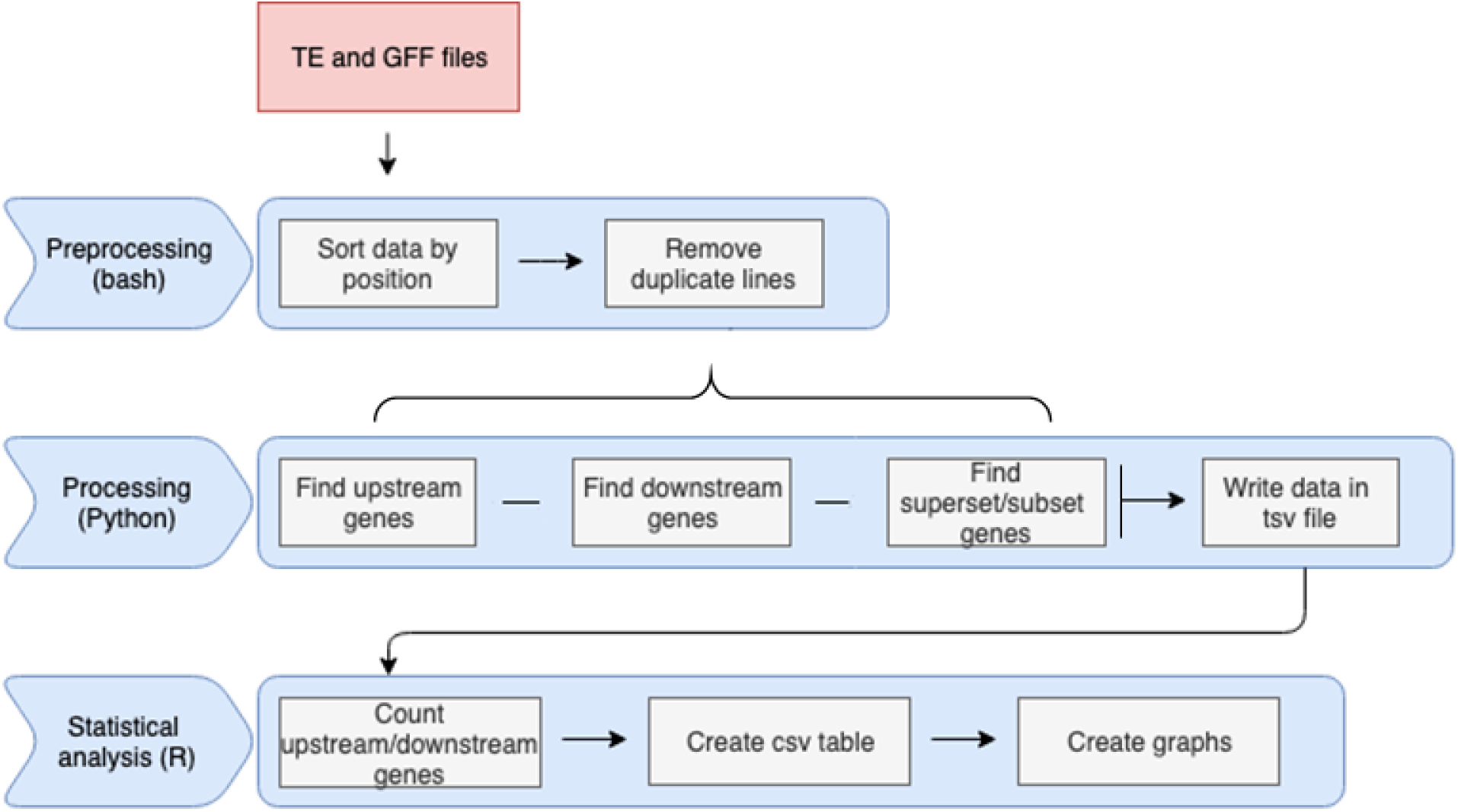
The workflow of the pipeline comprises three different steps: preprocessing, processing and statistical analysis.

### 2. Implementation

The first part of the algorithm was written using Python 3.7. The program contains six functions aiming to detect genes surrounding any TE, as well as other functions allowing to read the input data and to write the output in a TSV file. Here is a summary of those functions:

- check upstream gene: this function returns the closest gene to the TE positioned in an upstream location. Examples from Figure 1: Gene1-TE1; Gene4-TE5/TE6.
- check downstream gene: this function returns the closest gene to the TE positioned in a downstream location. Examples from Figure 1: Gene9-TE10; Gene11-TE11.
- check upstream overlap gene: this function returns genes with at least one base overlapping the upstream part of the TE. Example from Figure 1: Gene1-TE1.
- check downstream overlap gene: this function returns genes with at least one base overlapping the downstream part of the TE. Example from Figure 1: Gene9-TE10.
- check subset superset gene: this function searches for genes, which are either a subset (Gene7-TE9) or a superset (Gene6-TE7/TE8) of the TE.
- calculate distance: this function allows to calculate the distance between a TE and its closest gene.
- write data: this function writes an output file containing all the information about the genes detected by the previous functions.

The python output is a TSV file containing the TE-gene relationship information based on the physical structure of the genome as descripted above. For each case, the start position, end position, ID and strand information are recorded. The TE ID, type, strand and position are also reported. Graphical visualization of the data is obtained by R scripts with the 4.0.2 version. Using the output file from the Python script, descriptive statistics of the relation between TEs and genes in the genome are presented. The output of the R scripts is graphs and CSV files containing the values counted by the algorithm. There are three R scripts allowing users to report three different statistics:

- TE statistics, which show how many TEs and what types of TEs are present in the file and the distribution of TEs between those types.
- Overlap statistics, which show how many TEs have an overlap with genes, both upstream and downstream.
- Distance statistics, which show the number of TEs with an upstream or downstream gene within 0-500 bp, 500-1000 bp, 1000-2000 bp and more than 2000 bp intervals.

The genes reported as upstream or downstream of TEs in R scripts were defined based on the TEs’ strand as described in Figure 3.

**Figure 3.**
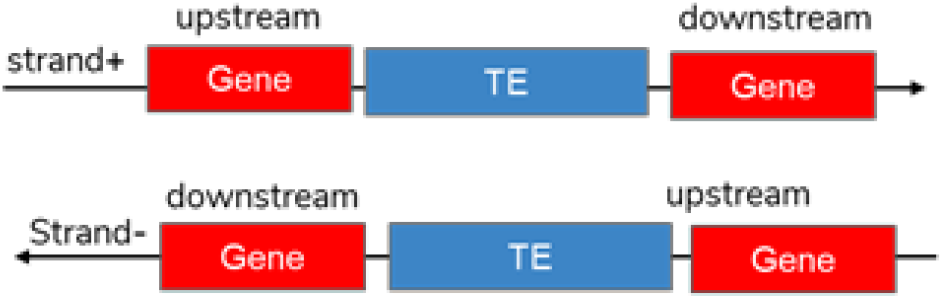
Representation of downstream and upstream genes based on autonomous TEs’ strand.

### 3. Operation

Our workflow requires R 4.0.2 or upper, Python 3.7 and can be run on any operating system with common specifications (1Go disk space, 4Go RAM, multicore CPU is recommended).

### Use case

In order to explain how the program works, we will use two example files named gene_testing_data and transposon_testing_data, which gather all possible scenarios of the position of TEs relative to their closest genes. These two input files can be downloaded from the github page. The following command must be run in order to execute the program:

~~~
python3 Multiprocessing/Create_Data_multipro.py \
        -g data/Gene_testing_data.tsv \
        -te data/Transposon_testing_data.tsv \
        -o result/output_TE.tsv
~~~

The -g argument is used to provide the gene file, the -te is used to come up with the TE file and the -o to supply the name of the result file. The output file is in TSV format. Once this file has been obtained, the R analysis can be run by adapting the following script line:

~~~
Rscript Rscript/number_te.r \
        -f result/output_TE.tsv \
        -o result/count_TE_transposons.pdf
Rscript Rscript/Overlap_counting.r \
        -f result/output_TE.tsv \
        -p result/overlap_TE_results.pdf \
        -o result/overlap_TE_results.csv
Rscript Rscript/Distance_counting.r \
        -f result/output_TE.tsv \
        -p result/distance_TE_results.pdf \
        -o result/distance_TE_results.csv
~~~

The R scripts will provide output files with the statistics regarding overlapping genes and LTR retrotransposons as shown in Figure 4, or distance between LTR retrotransposons and genes as shown in Figure 5, in table format as well as graph visualization.

**Figure 4.**
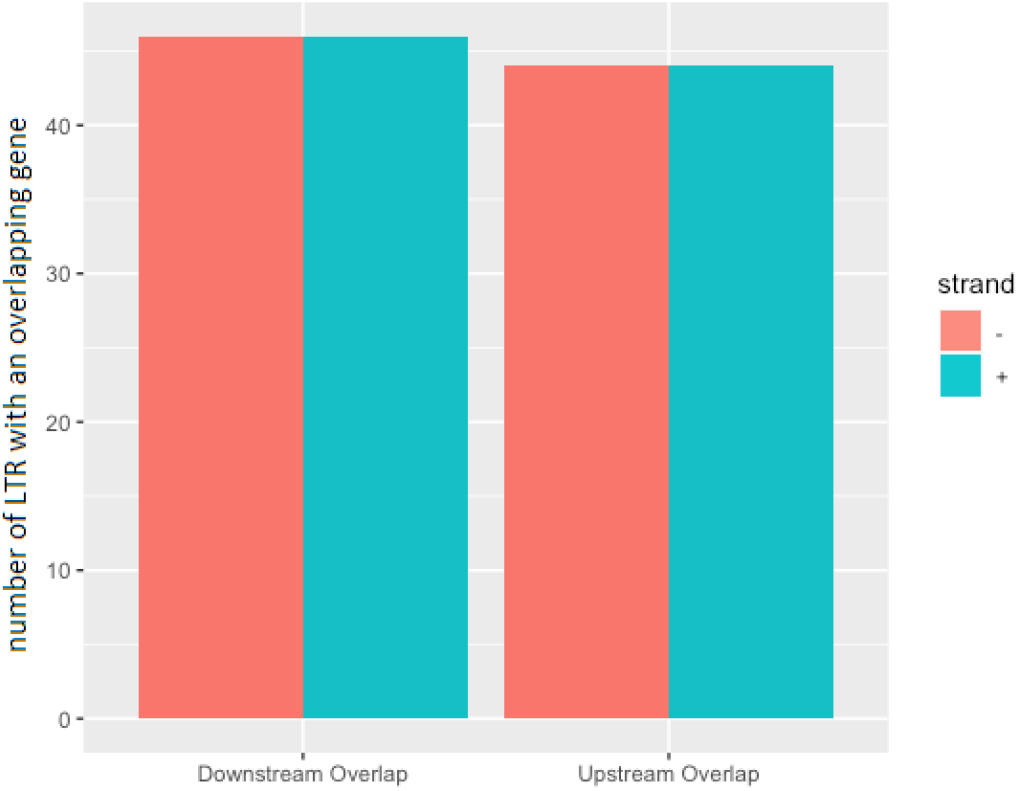
Number of sense LTR retransposons (strand+) and antisense LTR retransposons (strand-) with a downstream-overlap gene and upstream-overlap gene in *Prunus mandshurica*.

**Figure 5.**
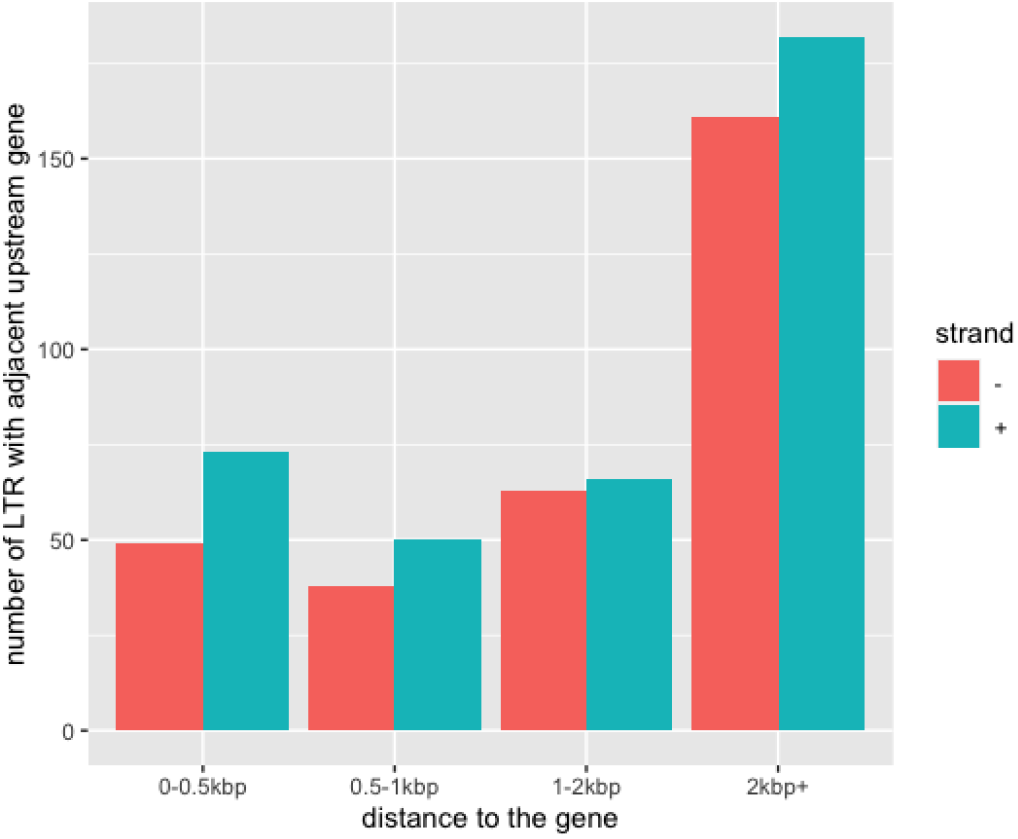
Number of sense LTR retransposons (strand+) and antisense LTR retransposons (strand-) with an adjacent upstream gene in *Prunus mandshurica*.

## Conclusion

Transposable elements are repetitive DNA sequences that have the ability to move within the genome. These mobile elements can play an important role in gene regulation and have a large impact on genome evolution. We have developed the first pipeline, which can report directly the relationship between TEs and its nearest genes among the genome. The accuracy of the tool has been verified by using a test dataset and running on two different TE annotation softwares. This pipeline could be useful to subsequently analyze potential effects of TEs on gene expression as well as on specific gene function.

## Software and data accessibility

Up-to-date source code and tutorials are available at: https://github.com/marieBvr/TEs_genes_relationship_pipeline. Archived source code as at time of publication are available from: https://doi.org/10.5281/zenodo.4442377.

The raw data of the Apricot genome, *Prunus mandshurica*, are available on the European Nucleotide Archive (ENA) under the project name PRJEB42606: https://www.ebi.ac.uk/ena/browser/view/PRJEB42606 and the assembled genome is available at https://www.rosaceae.org/node/10811682.

## Acknowledgements

We thank the INRAE BFP and the University of Bordeaux for the administrative support. Version 4 of this preprint has been peer-reviewed and recommended by Peer Community In Genomics (https://doi.org/10.24072/pci.genomics.100010).

